# Metabolomic, proteomic and single cell proteomic analysis of cancer cells treated with the KRAS^G12D^ inhibitor MRTX1133

**DOI:** 10.1101/2023.03.23.533981

**Authors:** Benjamin C. Orsburn

**Affiliations:** The Department of Pharmacology and Molecular Sciences; The Johns Hopkins University School of Medicine, Baltimore, MD, USA, 21205

**Keywords:** Single cell proteomics, metabolomics, proteomics, KRAS, MRTX1133

## Abstract

Mutations in KRAS are common drivers of human cancers and are often those with the poorest overall prognosis for patients. A recently developed compound, MRTX1133, has shown promise in inhibiting the activity of KRAS^G12D^ mutant proteins, one of the main drivers in pancreatic cancer. To better understand the mechanism of action of this compound I performed both proteomics and metabolomics on four KRAS^G12D^ mutant pancreatic cancer cell lines. To obtain increased granularity in the proteomic observations, single cell proteomics was successfully performed on two of these lines. Following quality filtering, a total of 1,498 single cells were analyzed. From these cells 3,140 total proteins were identified with approximately 953 proteins quantified per cell. At 48 hours of treatment, two distinct populations of cells can be observed based on the level of effectiveness of the drug in decreasing total abundance of the KRAS protein in each respective cell, results that are effectively masked in the bulk cell analysis. All mass spectrometry data and processed results are publicly available at the www.massive.ucsd.edu at accessions PXD039597, PXD039601 and PXD039600.

**Abstract Graphic:** 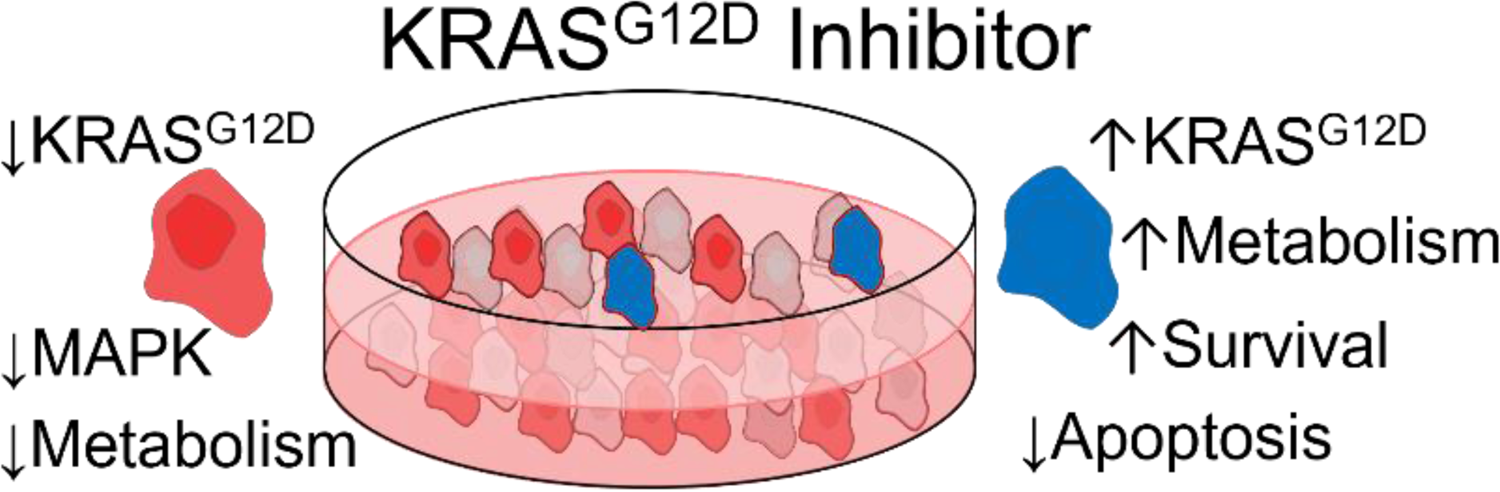

## Introduction

KRAS mutations are found in up to 25% of solid tumors, and typically those with the worst prognosis.^1,2^ Pancreatic cancer is the best example of this, as a cancer with a dismal survival rate of only 44% after 5 years. Mutation in KRAS, and specifically the KRAS^G12D^ mutation, are particularly prevalent in pancreatic cancer. A recently described compound, MRTX1133, builds on recent successes with KRAS small molecule inhibitors, as the first inhibitor described for the KRAS^G12D^proteoform.^3^ MRTX1133 differs from successful KRAS^G12C^ inhibitors in that it is purported to bind through noncovalent mechanisms.^4,5^ To better evaluate the mechanism of action of this exciting new inhibitor, I treated four human pancreatic cancer cell lines with the drug for 48 hours and performed standard label free proteomics and global untargeted metabolomics.

Work in our lab recently described the use of single cell proteomics (SCP) to better understand the heterogeneity of response in the proteomes of cells treated with sotorasib, a KRAS^G12C^ inhibitor. While more of a proof of concept, the work led to the development of tools for multiplexed SCP using a TIMSTOF instrument. Multiple limitations existed in the study, including the relatively small number of cells that were analyzed. We were, however, able to recapitulate many of the findings of both bulk proteomic and single cell transcriptomic studies of drug activity, due largely to profound effects sotorasib had on cell cycle. Despite limitations, we were able to highlight new findings that were not previously revealed by other approaches.^6^ Following optimization to alleviate bottlenecks on the sample preparation and data analysis side, I have applied the resulting methods to two cancer cell lines treated with MRTX1133.

## Methods

### Cell culture and drug treatment

All cell lines were obtained from ATCC between March and December of 2022 and were cultured according to vendor instructions. PANC 0813, PANC 0304 and PANC 0203 were grown in RPMI 1640 (ATCC 30-2001) supplemented with 15% fetal bovine serum (ATCC 30-2020) and 10 units of human insulin (Fisher). ASCP-1 was grown in RPMI 1640 with 10% FBS. All cell culture media contained 10 mg/mL Penn Strep antibiotic solution (ATCC 30-2300). All cell lines were passaged a minimum of 3 times prior to treatment with 10 nanomolar MRTX1133 for 48 hours. Cells were harvested by vacuum aspiration of the cell culture media. The adherent cells were briefly rinsed in 3 mL of 0.05% Trypsin plus EDTA solution (ATCC 30-2001). This solution was rapidly aspirated off and replaced with 3 mL of the same solution. The cells were examined by light field microscopy and incubated at 37⁰C with multiple examinations until the adherent cells had lifted off the plate surface. The active trypsin was then quenched by the addition of 7 mL of the original culture media. The 10 mL solution was transferred to sterile 15 mL Falcon tubes (Fisher) and centrifuged at 300 *x g* for 3 minutes to pellet the cells. The supernatant was gently aspirated off and the cells were resuspended in PBS solution without calcium or magnesium with 0.1% BSA (both, Fisher Scientific) at 1 million cells per mL as estimated by bright field microscopy. Approximately 2 million cells were taken for bulk proteomic and metabolomic analysis. Cells for single cell aliquoting were gently dissociated from clumps by slowly pipetting a solution of approximately 1 million cells through a Falcon cell strainer (Fisher, 353420) and the cells were placed on wet ice and immediately transported across the street to the JHU Public Health sorting core. Non-viable cells were labeled with a propidium iodide solution provided by the core facility and briefly vortexed prior to cell isolation and aliquoting.

### Global Metabolomics

A solution containing approximately one million cells from each condition was centrifuged at 13,000 *x g* for 15 seconds at speed to obtain a solid pellet. The supernatant was aspirated off and replaced with 300 microliters of 70% LCMS methanol in LCMS water solution. Pellets were resuspended with vigorous vortexing and placed at −80⁰C for approximately 48 hours. Following storage, the cells and methanol solution were thawed on wet ice and resuspended with vigorous vortexing, followed by centrifugation at 13,000 *x g* for 5 minutes to precipitate proteins and cellular debris. The top 250 microliters of supernatant were removed to limit disturbing of the lower material and was dried to completeness via SpeedVac at 25⁰C. The dried metabolite extract was resuspended in 50 microliters of 5% LCMS grade acetonitrile 0.1% formic acid in LCMS grade water. A pooled sample containing equivalent amounts of all control and treated metabolite extracts was prepared for use as the reference sample for chromatographic alignment. Two microliters of solution were used for each of four replicate LCMS injections. For LCMS analysis a Q Exactive “Classic” system coupled to Dionex U3000 UHPLC was used. Separation was performed on a 2.1 mm x 15 cm HyperSil Gold Colum with 2 micron particle size and a flow rate of 300 microliters/minute for the active gradient. The LCMS method parameters have been deposited at www.LCMSmethods.org as “Q Exactive Positive Metabolomics” in the 2019 method release. Briefly, MS1 scans were acquired from 90-850 m/z at 70,000 resolution, with an AGC target of 3e6 charges or a maximum injection time of 50 ms. Data dependent acquisition with a loop count of 3 was used to fragment ions which were accumulated with a target of 2e5 charges or a maximum injection time of 50 ms. A three step collision energy was used with 10, 20, and 60 CE used for each fragment ion. Source conditions were determined through the automated method with Exactive Tune 2.11 SP1 for this flow rate. All resulting output files were processed in Compound Discoverer 3.1 using a combination of 3 human metabolite libraries from mzCloud, ChemSpider, and a local desktop library of 4,000 human metabolites provided by Thermo Scientific. Compound identities were prioritized in the case of conflict between databases in the order described above.

### Bulk proteomics

A solution containing approximately one million cells from each condition was rapidly centrifuged at 13,000 *x g* for 60 seconds at speed to obtain a solid pellet. The sorting solution was carefully aspirated off and the cell pellet was resuspended with 200 microliters of S-Trap lysis buffer (5% SDS in 100mM TEAB, ProtiFi). The pellet was suspended with rigorous vortexing prior to being placed in an ultrasonic water bath for 15 minutes at 37⁰C for further cellular lysis. Fifty microliters of each sample were taken for reduction at 95⁰C for 5 minutes in 20mM DTT followed by benchtop cooling to room temperature and alkylation with 30mM iodoacetamide for 20 minutes at room temperature in a light tight drawer. Pierce “Easy Aliquot” single use reagents were used for both reduction and alkylation. Another 50 microliters of each solution were taken for digestion without reduction and alkylation for the later construction of bulk cell digest carrier channels. Digestion in both cases was performed using an S-Trap mini (Protifi) spin column according to vendor instructions with an approximate 200 micrograms of total protein load and a 2 hour digestion at 47⁰C. Peptides were eluted following digestion and vacuum centrifuged to dryness and resuspended in 0.1% formic acid. Peptides were quantified using a peptide colorimetric assay (Pierce 23290).

All proteomic analysis was performed on a TIMSTOF Flex mass analyzer (Bruker) coupled to an EasyNLC 1200 system (Proxeon) by a 15 cm x 75 micron PepSep column with 1.9 micron Reprosil particles. The data acquisition method was the default diaPASEF method “short gradient diaPASEF” included in TIMS Control 4.0 2023. The chromatography gradient ramped from 8% buffer B (80% acetonitrile in 0.1% formic acid) to 35% B in 25 minutes with a flow rate of 350nL/min prior to a rapid increase to 100% B at 500 nL/min by 30 minutes. Baseline conditions were restored at the beginning of each chromatography gradient by the HPLC prior to loading the next sample.

### Single cell aliquoting

Single cell proteomics sample preparation, analysis and data processing were performed as described previously.^6^ Briefly, single cells were aliquoted using an analog MoFlo sorter into cold 96 well plates containing 2 microliters of LCMS grade acetonitrile. At the completion of each plate aliquoting they were immediately sealed and placed in an insulated box of dry ice with the wells pressed into the material to ensure rapid cooling. After approximately 5 minutes, each plate was transferred to a cooler of wet ice for transport back to our lab and −80⁰C storage. Estimations of cellular viability and cell aliquoting efficiency were provided by the core director who has over 30 years of cell sorting and aliquoting experience and significant experience in the analysis of pancreatic cancer cell lines by flow cytometry.^7–9^ Digital reports of FACs data was provided by email following the isolation procedure and are available upon request.

### Single cell lysis, digestion and combination of multiplexed cells

Single cells were placed into storage followed extensive use of tape to help ensure plate seal integrity during storage. A full protocol describing all steps of cell lysis, digestion and TMT labeling has been permanently published at Protocols.io and can be accessed at dx.doi.org/10.17504/protocols.io.yxmvm2w16g3p/v1. Cell lysis was further driven to completeness and acetonitrile was removed by placing each plate directly from cold storage onto a 95⁰C hot plate for 90 seconds. The plates were cooled to room temperature on a benchtop with static free surface covers. The author performed all aliquoting of reagents while standing on a static free floor mat at all stages of the preparation. Periodically a Zero Stat “static gun” clearly labeled in fluorescent pink tape “NOT A GUN” to help ensure author survival during late night sample preparation was used to further minimize the negative side effects of static electricity on single cells (Fisher Scientific, NC9663078). Dried lysed cellular lysate was digested using a solution of 5 nanogram/microliter LCMS grade trypsin (Pierce) in 0.1% n-Dodecyl-beta-Maltoside Detergent (DDM, Thermo Fisher, 89902) and 50mM TEAB. One microliter of trypsin solution was used for all single cells and blank wells and four microliters were used for digestion of the carrier channel lanes. The excess trypsin was used as an extension of the “sacrificial carrier” concept recently described in Choi *et al*.,.^10^ The plates were tightly sealed and incubated for 2 hours at 45⁰C in an Orbital shaker with centrifugation every 30 minutes to condense evaporate. Following digestion, the plates were cooled to room temperature and centrifuged again to precipitate all liquid solution. TMT Pro 18 reagents previously prepared and aliquoted at a concentration of 20 nanograms per microliter were used to label all wells and sacrificial trypsin peptides by adding 1 microliter and incubating for 30 minutes at room temperature with orbital shaking. A solution of 0.5% hydroxylamine in TEAB was used to quench remaining TMT by adding 0.5 microliters to each well with a 20 minute incubation at room temperature with orbital shaking. Each plate was then dried by SpeedVac at room temperature, which took approximately 5 minutes for each pair of plates. Ten unit resolution TMT channels from the TMTPro 18 kits (Thermo Fisher) were used. The method blank was labeled with the 126 reagent in all comparisons and 135n was used for the carrier. To minimize the effects of isotopic impurity carryover, staggered reagents were utilized from the kit, with 127n, 128c, 129n, 130c, 131n, 132c, 133n and 134c used to label single cell wells.^11^ TMT labeled single cells and carrier channels were combined in a staggered manner to ensure each TMT plex contained a mixture of the control and treated carrier channel digest and both control and treated cells. For example, injection well A1 on the autosampler would contain control cells as 127n, 128c, 129n, and 130c, with labeled treated cells occupying the remainder of the multiplex set. Autosampler well A2 would contain the reversed set, with the first four channels containing treated cells, with control cells as the last four channels. The net result is that each well in the autosampler plate contains a carrier channel that is an equal mixture of control and treated cells. In addition, four control cells and four treated cells were measured in each LCMS experiment. By alternating loading of single cell plates, we can obtain a pseudo-random injection pattern where a different set of single cells from each batch with alternating labels are injected consecutively. The cell identities were deconvoluted during the final data analysis in Proteome Discoverer.

### Comparison of FACs sorted and bulk cell homogenate carrier channels

To compare the two competing methods for carrier channel preparation, aliquots from bulk cell lysates that were not reduced and alkylated as mentioned above were labeled with the 135n channel according to manufacturer’s protocol, with the exception that a 4:1 ratio of label to peptide was utilized. Pre-labeling peptide concentrations were used to estimate the post-labeled concentrations. Control and treated lysates from PANC 0813 and ASPC-1, respectively, were combined to create a pooled carrier lysate for each cell line. An estimated 40 nanogram mixture was used to resuspend each single cell set using the same combination method described above for the FACs sorted single cell lysates. The end goal was an equivalent pooled carrier channel and both control and treated single cells analyzed in each LCMS injection with the channels flipped in each respective injection.

### Single cell data analysis

As previously described,^12^ I utilize a single point recalibration method using the 135n carrier channel signal to adjust the reporter ion region. This secondary calibration allows tighter mass accuracy tolerances to be used during the final data analysis, resulting in reduced background noise. For all ASPC-1 and PANC 0813 cells to meet the QC requirements for inclusion in this manuscript, the linear calibration shift applied was +0.0141 Da. This was performed using an updated version of the pasefRiQ calibrator with a GUI interface and the ability to batch recalibrate MGF files. This software is openly available and permanently published at DOI: 10.5281/zenodo.7259511. The recalibrated output files were processed in Proteome Discoverer 2.4SP1 using SequestHT and Percolator. Briefly, the MS/MS spectra were binned into 100 Da segments and filtered so that only the top 12 most abundant ions from each bin were retained. The resulting cleaned spectra were searched with a 15ppm MS1 tolerance and 0.03 Da MS/MS tolerance. TMTPro labels were considered static on the N-terminus and dynamic on lysines to allow for the search for lysine acetylation, methylations and dimethylation. Methionine oxidation was the only additional dynamic modification. The default cutoffs for Percolator PSM validation and peptide and protein FDR determination were employed in all analyses. Reporter ions were integrated using a 35ppm mass tolerance window (17.5 ppm up and down) and quantification values were only used for unique peptides. Multiple consensus workflows were performed on the resulting MSF files to assess different normalization methods. The final data report for downstream analysis used a sum based normalization of all PSM signal for TMTPro channels 127-133 as well as the raw non-normalized abundance values. The 134 single cell channel was discarded in all runs due to apparent inflation from impurities in the 135 reagent channel.

### Uniform Manifold Approximation and Projection (UMAP)

The normalized output sheet for all single cells from Proteome Discoverer was converted to .CSV and loaded into Perseus 1.6.17.^13^ Perseus was previously set up to communicate with R 4.1.3 as described previously.^14^ All single cell channels were added into Perseus as “Main” data points with the Protein Accession and Description added as “Text” variables. All zero datapoints were replaced with normal distributions with each single cell used as the distribution matrix individually. The cells were then grouped under row categories describing their individual cell and dose conditions. UMAPs were generated using the default settings for Euclidean distribution with 15 neighbors, 2 components, random state of 1 and a minimum distance of 0.1. The output was plotted using the multi-scatter node in Perseus.

### Clustering Drug Treated Cells by Relative KRAS^G12D^ Expression

The raw non-normalized abundance values for the KRAS protein for each cell with a measured abundance 3 times the calculated noise level was exported from the Proteome Discoverer output for each control and MRTX1133 treated cell. The output text file was analyzed in GraphPad Prism to determine the median and mean protein abundance. MRTX1133 treated cells with an abundance greater than the mean for all treated cells were manually flagged for clustering. Twenty five PANC0813 cells with a level of KRAS expression greater than the treated mean were analyzed in the SimpliFi platform as a single group compared to 107 cells with lower levels of relative expression as the control group. Analysis was by default parameters with the exception that missing values were allowed to be up to 50% per protein to account for the missing values inherent in single cell proteomics.

### Data Availability

All proteomics results have been deposited and can be viewed in the SimpliFi cloud platform. The cell lines with a single and double copy of the KRAS^G12D^ mutant form were processed separately for ease of visualization. The PANC 0813 and ASPC-1 proteomics data can be found at the following link: https://simplifi.protifi.com/#/p/4fd12e70-9809-11ed-b0f0-03f62ebcf0cd. PANC 0203 and PANC 0403 can be found here: https://simplifi.protifi.com/#/p/28a07700-b93a-11ed-92f4-b7d04df347c4. The original RAW and processed files are divided between 3 openly available repositories at www.Massive.UCSD.edu: PXD039597, PXD039601 and PXD039600.

## Results and Discussion

### MRTX1133 treatment alters central metabolism

Global reversed phase positive metabolomics identified 940 distinct small molecules after standard adduct reduction processes. Of these, 781 could be assigned a putative molecular formula and 264 could be assigned a putative name within the cutoffs uses for this study (**Supplemental File 1**). The most striking changes were that drug treatment revealed a universal depletion of ATP precursors in all cell lines following MRTX1133 treatment. Notably, adenosine (**Figure 1A**), AMP (**Figure 1B**), and ADP were observed at reduced relative concentrations. NAD precursors were likewise reduced by drug treatment in all four cell lines. In contrast, many amino acids such as tyrosine and glutamate (**Figure 1C**) were observed at increased relative concentration in drug treated cells. A smaller number of central metabolites appeared to demonstrate mutant copy number specific effects. PANC 0203 and 0403, which have single copies of the mutant gene appear to have increased levels of glutamine following treatment, while PANC 0813 and ASPC1 which have two copies of the protein, appear to have decreased levels of this metabolite following treatment (**Figure 1D**). While this is a small group for comparison, the number of KRAS mutant gene copies numbers have been previously linked to metabolic reprogramming effects.^15^

**Figure 1.**
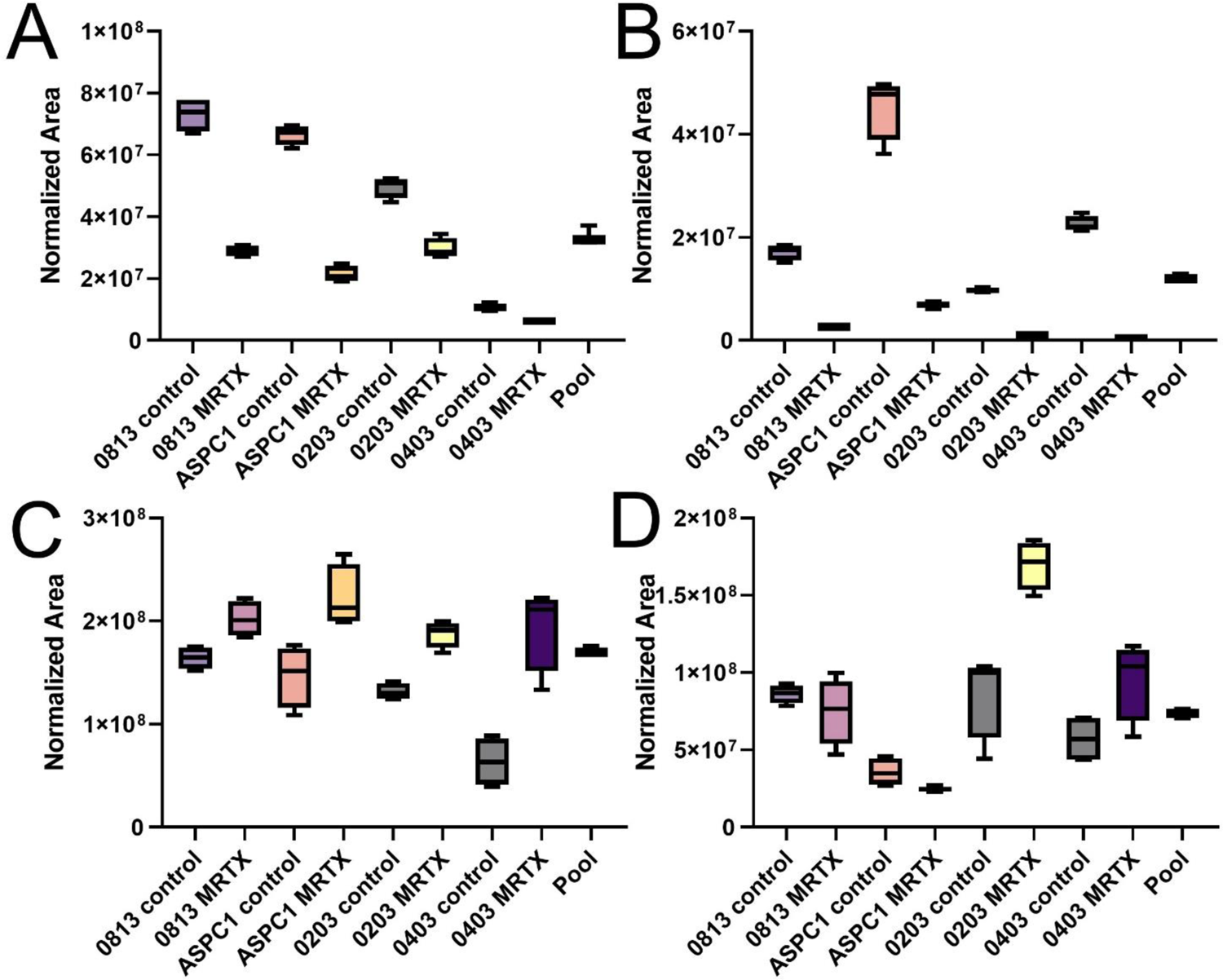
Representation of global metabolomic alterations following MRTX1133 treatment. **A**. Adenosine levels are decreased in all cell lines following treatment. **B.** The same trend is observed in AMP. **C**. Drug treatment leads to increased intracellular glutamate in all cell lines. **D.** Glutamine levels following treatment suggest a different effect in the cell lines harboring single or double copies of the mutant gene.

### Proteomic bulk cell homogenate data

The bulk cell proteomics data largely produced expected changes in all cell lines. MRTX1133 acute treatment resulted in a consistent decrease in abundance in the central MAPK pathway. ERK (MAPK3) was reduced in all cell lines following treatment, as previously reported^4^ as was MAPK1 (**Supplementary** Figure 1). SimpliFi pathway analysis identified the Rho GTPase pathway as the seventh most significantly altered pathway between control and treated lines (R-HAS-563220; 2.04 × 10^-60^). Similarly, MAPK was the 9^th^ most significant altered pathway (R-HSA-5687128; 7.41 × 10^-53^). The MHC Class II antigen presentation system fell (R-HSA-2132295; 9.03 × 10^-60^ between these two pathways which is also expected based on work demonstrating high levels of MHC level expression in KRAS mutant cells.^16^

### Single cell proteomics summary

We have recently described the use of a TIMSTOF mass spectrometer for multiplexed SCP. While we determined that higher loading channel concentrations could be utilized with little decrease in quantitative accuracy, they came at the expense of increased missing values.^12^ In this study I employed a protocol more similar to the original SCoPE-MS study^17^ in which 200 single cells from each condition were used as the carrier channel in most experiments. For a smaller subset of cells, a pooled diluted homogenate was used as the carrier channel. By mixing the loading channels from each experimental condition and using a pseudo-random mixing and loading protocol, I can combine SCP samples from multiple conditions within each LCMS experiment. While this may seem to be an obvious way to set up a study of this sort, this is not yet possible in commercial products such as the ProteoChip kit designed for use on the CellenOne systems.^18,19^ At 200 cell carrier with an estimate of 200 picograms of protein per cell the starting protein load for each injection should be approximately 41.6 nanograms. In the two cell lines described 953 proteins were quantified, on average, when using a single search engine. In total, 3,143 protein groups were identified. We have found that the use of multiple search engines leads to more than a twenty percent increase in identifications over a single search engine alone. Similar observations have been recently reported by others.^20^ In addition, multiple advanced tools for single cell analysis such as isobaric tag enabled match between runs ^21^, and posterior error adjustment can lead to a further increase in total identification rates.^22^ The complete report of all proteins and quantification values for all single cells is provided as **Supplemental File 2.**

### Protein post translational modifications can be identified in nearly all single cells

As previously reported for the H358 cell line, I can detect and quantify multiple classes of protein PTMs in single cells in this study. The most abundant and readily observed PTMs are histone modifications, particularly the well characterized and tryptically convenient K28 and K80 lysine modification sites. Bar plots demonstrating the abundance of the K28 methylated and dimethylated peptides are shown in Figure 2. The former is commonly observed, with signal in 75.8% (440/580) of the PANC 0813 single cells shown. When considering the possibility of methylation or dimethylation occupying the K28 site, 98.7% of single 0813 cells (573/580) demonstrate quantifiable signal. These results are consistent for both cell lines in all experiments (data not shown). The ability to detect protein PTMs is directly related to both the protein total copy number as well as the total sequence coverage obtained for the protein. The tryptic peptide containing the K28 modification is identical in at least 3 different human histone proteins annotated in SwissProt UniProt database today, all of which exist in excess of one million copies in most mammalian cells.^23^ As such, it is difficult to interpret the meaning of these PTMs in these data, as each signal observed may be a composite of modifications from multiple original histone proteoforms. These results suggest that the analysis of histone PTMs in single human cells would be a practical avenue for research. However, as in traditional proteomics, the use of alternative proteases^24^ or lysine derivatization prior to tryptic digestion^25^ would be required for accurate assignment of the PTM to the originating proteoform.

**Figure 2.**
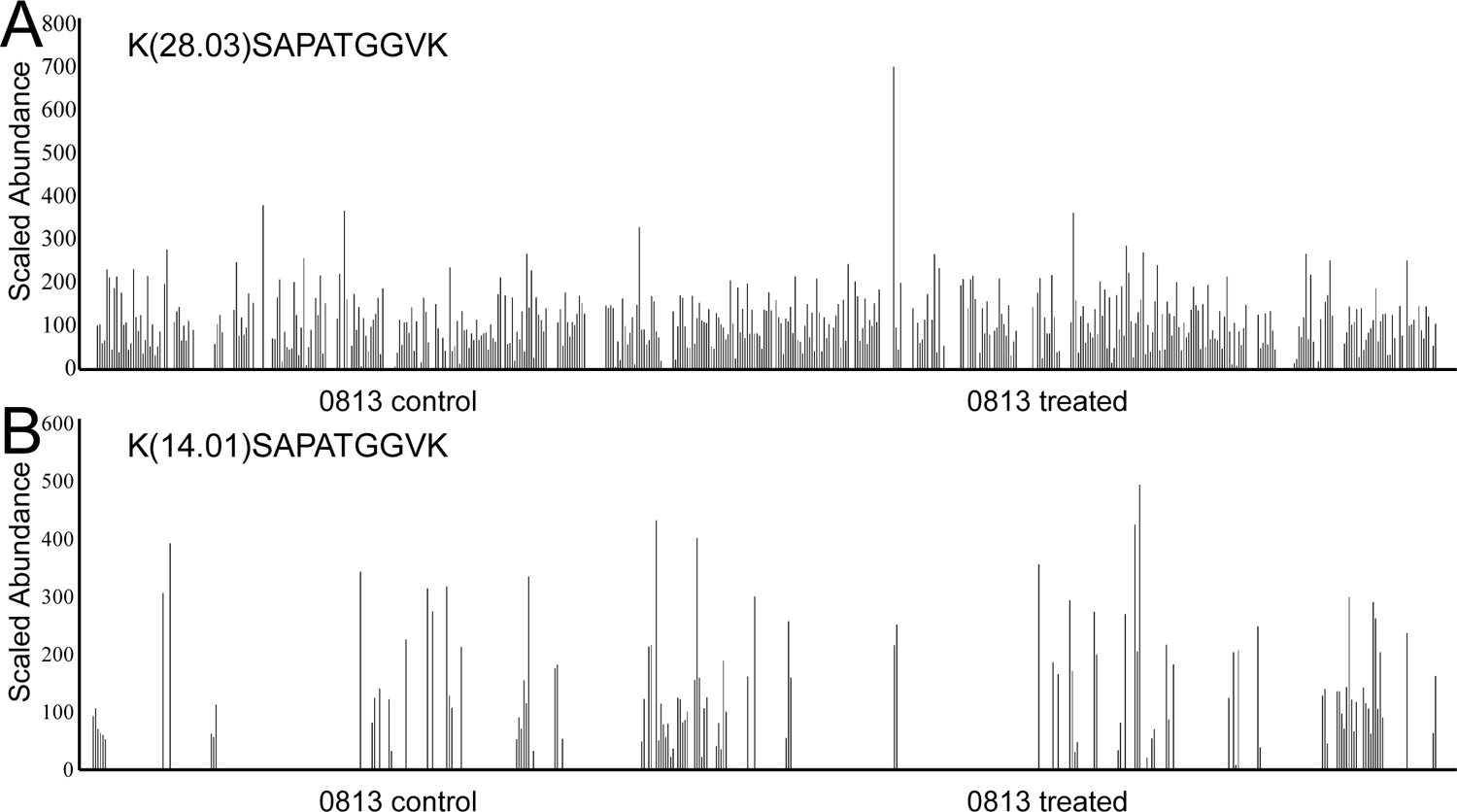
The measured abundance of two PTMs on histone 3 with 580 single 0813 cells shown. **A.** The K28 dimethylated peptide was observed in over 75.8% of these cells. **B.** A single dimethylation on this lysine was observed less frequently.

### Unsupervised analysis can discriminate between similar pancreatic cancer cell lines

One quality control metric for a single cell proteomics methods that has been employed by multiple groups is the demonstration of cell type clustering by unsupervised statistics such as PCA.^18,26,27^ If HeLa cells and HEK293 cells form distinct clusters when analyzed together, it stands to reason that the method is sound. One criticism of this method is that there are considerable differences between the size and protein content of these two common laboratory cancer cell line strains. ASPC-1 and PANC 0813 are two highly similar pancreatic cancer cell lines that were derived from patient samples using the same immortalization stategy.^28^ In addition, both cell lines possess two copies of the KRAS^G12D^ mutant gene. One major difference is that the ATCC recommends supplementing media for growing PANC 0813 with insulin, where none is recommended for ASPC-1. In limited experiments in our lab, we found this addition to be critical for cell survival (data not shown). As shown in Figure 3, a principal component analysis can resolve these two cancer cell lines. A recent preprint demonstrated that the largest proteomic effects in single cell measurements are simply cell size and recommended the normalization of single cell signal using histones as the normalization factor.^29^ While normalization with Histone H4 does appear to reduce the gap in clustering between these two cell lines, it did not lead to a coalescence of these two groups (data not shown). It is worth noting that previous studies have demonstrated large proteomic effects from insulin stimulation in the media of cancer cell lines.^30^ It is therefore possible that the true factor driving clustering visualized in this figure is insulin stimulation alone. However, this is still likely a more biologically meaningful metric for SCP method development than the unsupervised clustering of HeLa and HEK 293 cells.

**Figure 3.**
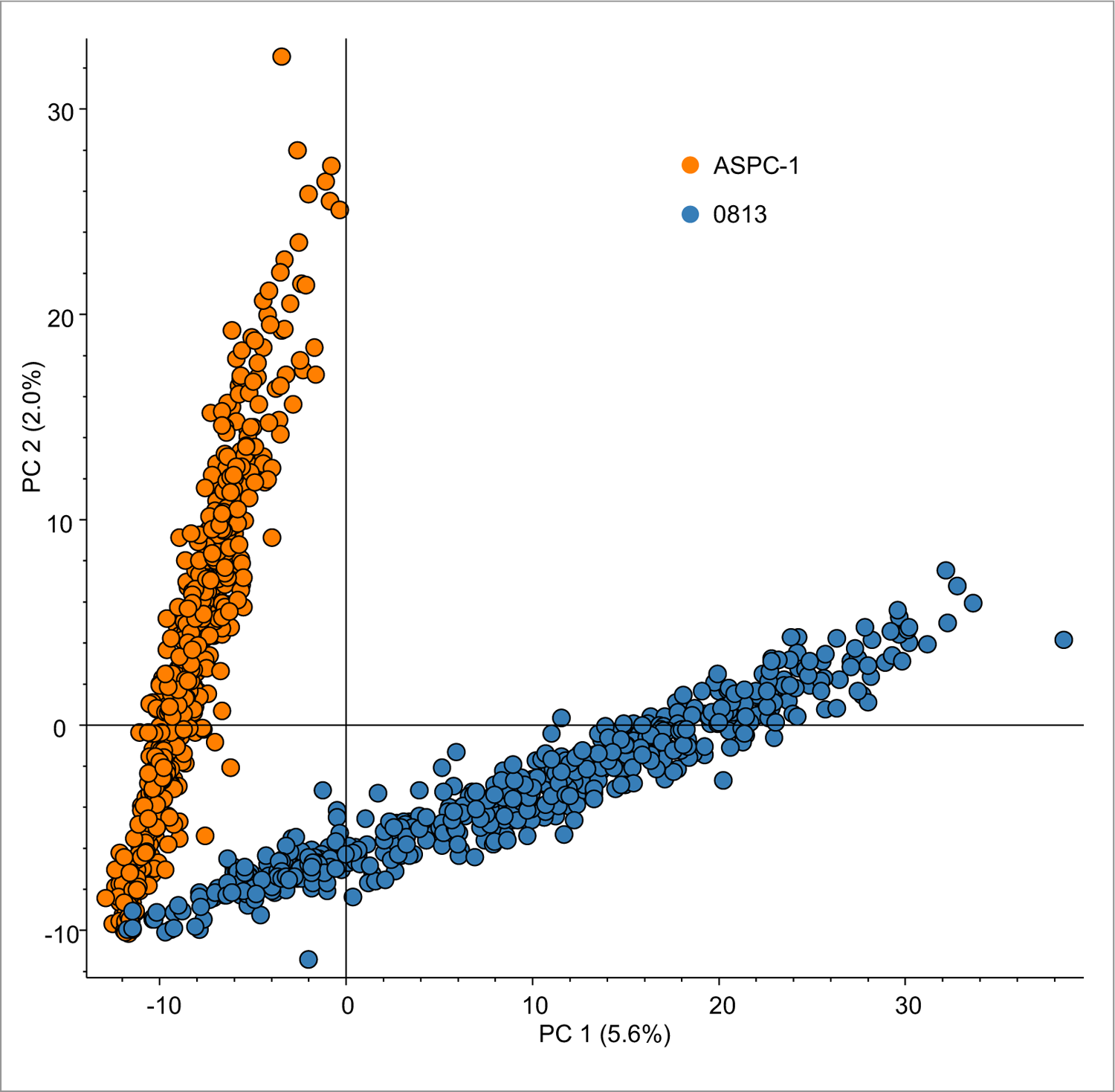
Principal component analysis of random injections of two similarly derived KRAS^G12D^ mutant pancreatic cancer cell lines (*n* = 1169).

### How carrier channels are prepared alters the proteins that can be detected in SCoPE-MS

There are two main strategies today for the preparation of carrier channels for multiplexed SCP. The original study by Budnik *et al*., utilized a collection of single cells which were lysed and prepared as the carrier channel.^17^ Studies utilizing liquid handling devices such as the CellenOne system, a miniaturized robotics system based on the SciFlex Arrayer^31^, likewise use this strategy.^18,32^ In Orsburn *et al*., we prepared a bulk cell homogenate of cells to be analyzed, labeled the homogenate with the 134c channel and prepared a dilution series to obtain an experimentally determined ideal carrier channel concentration for analysis.^6^ While our hypothesis was that this consistency in carrier composition would lead to a higher consistency between samples, an analysis of the two preparation methods appeared warranted. To evaluate the differences between these two approaches, I prepared carrier channels for PANC 0813 and ASPC-1 using both approaches. As shown in Figure 4 unsupervised analysis clusters the samples into four groups based on cell line and the carrier channel creation method.

**Figure 4.**
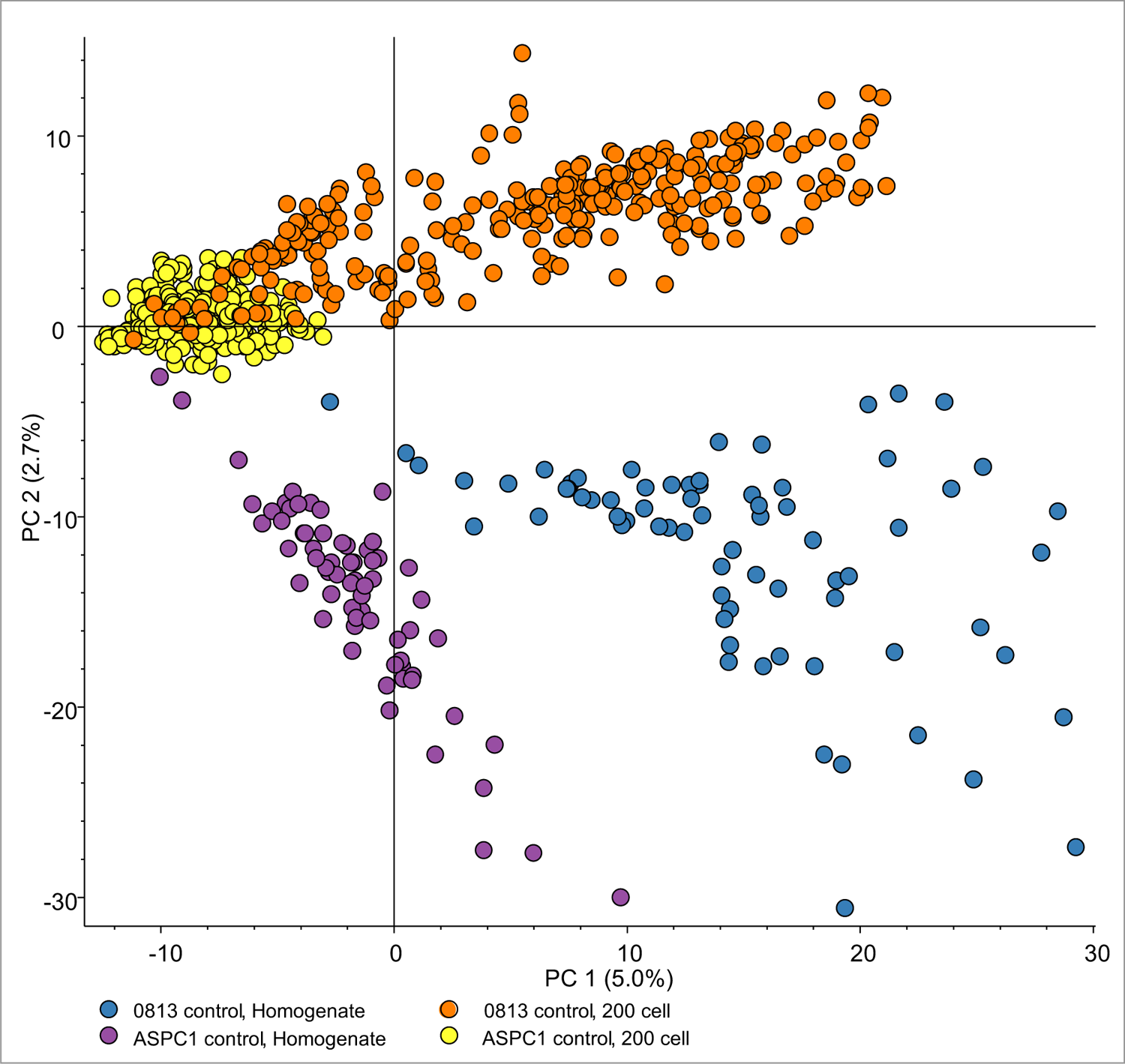
Principal component analysis of PANC 0813 and ASPC1 cells analyzed using either a bulk proteomics homogenate for a carrier channel, or a FACs sorted 200 cell carrier channel (*n* = 735).

**Figure 5.**
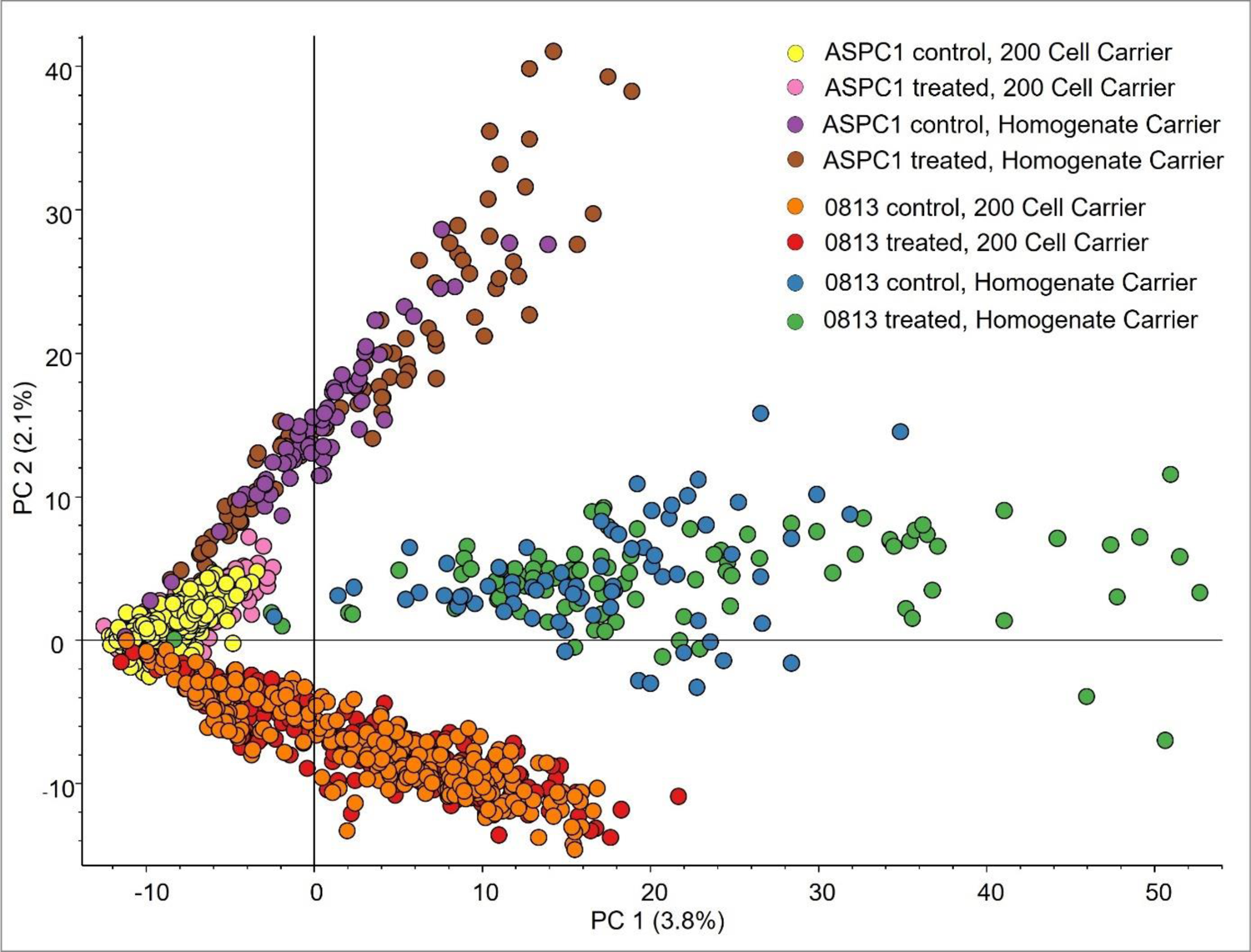
Principal component analysis of 1,498 KRAS^G12D^ control and MRTX1133 treated cells. Experiments where the carrier channel was prepared from FACs isolated single cells are denoted as “200 cell carrier”. Experiments where the carrier was prepared from a bulk cell homogenate that was labeled and diluted are denoted as “Homogenate Carrier”.

### Bulk cell lysate carriers increase the representation of proteins involved in apoptosis

To assess the differences imparted by the two carrier channel preparation strategies, I examined the functions of proteins exclusively identified in each carrier channel experiment using a simple plus/minus filter in Venny.^33^ Proteins unique to each carrier channel preparation method were analyzed for function in StringDB.^34^ The most striking difference between groups was the presence of apoptotic and programmed cell death proteins which appeared almost exclusive to files where carrier channels were prepared from bulk cell homogenates. Manual evaluation of the original proteomic data confirmed that proteins such as Caspase 6 and Caspase 8 were exclusive to these samples (**Supplemental** Figure 1).

### The single cell proteomics data largely recapitulates bulk proteomics data

As mentioned above, MRTX1133 treatment at this time point leads to a decrease in MAPK pathway protein levels, such as a 2.38-fold decrease in MAPK1 levels in PANC 0813 and a 3.46-fold decrease in the same in ASPC-1. As a 2-fold differential is a frequently used cutoff in preliminary proteomic data analysis, and this protein is known to be linked to drug treatment, an analysis of MAPK1 in PANC 0813 was a practical metric for SCP data quality.

During the first analysis of these data, it did not appear that the single cell proteomics recapitulated these findings. This was despite a relatively high detection rate for MAPK1 which was quantified in 61.4% of PANC 0813 cells in this study This led to the release of a preprint with a considerably pessimistic tone regarding the current state of single cell proteomics and this method of approach. Helpful discussions originating from the preprint manuscript as well as a complete reanalysis of these files presented at ASMS 2023 by Jim Palmieri (data to be published elsewhere), suggested flaws existed in the quantitative data analysis.

For a more direct analysis the abundance values for MAPK1 were plotted during each stage of data normalization. In PANC 0813, a mean fold change of −1.48 was observed when evaluating the raw unadjusted protein abundance measurements for MAPK1 in control versus treated cells. A student’s unpaired t-test found a significant difference between the control and treated cells (p = 0.031, n = 553). However, both sequential normalization and abundance scaling reduced both the observed ratios and apparent significance. Sum based normalization reduced the fold change to −1.11 at a p-value of 0.238 and scaling further reduced the median ratio to −1.09 at a p-value of 0.327, suggesting no change and no significance (Figure 6A). It should be noted that when we previously utilized a two proteome standard to determine the extent of the carrier channel effects on a TIMSTOF instrument, we observed that known ratios of 5:1 were compressed to approximately 3:1 when using a 200x carrier channel. Using a simple linear extrapolation for MAPK1, the −2.38-fold differential observed in bulk proteomics would represent a strikingly similar expectation of −1.50-fold for MAPK1 at this carrier level on this platform.

**Figure 6.**
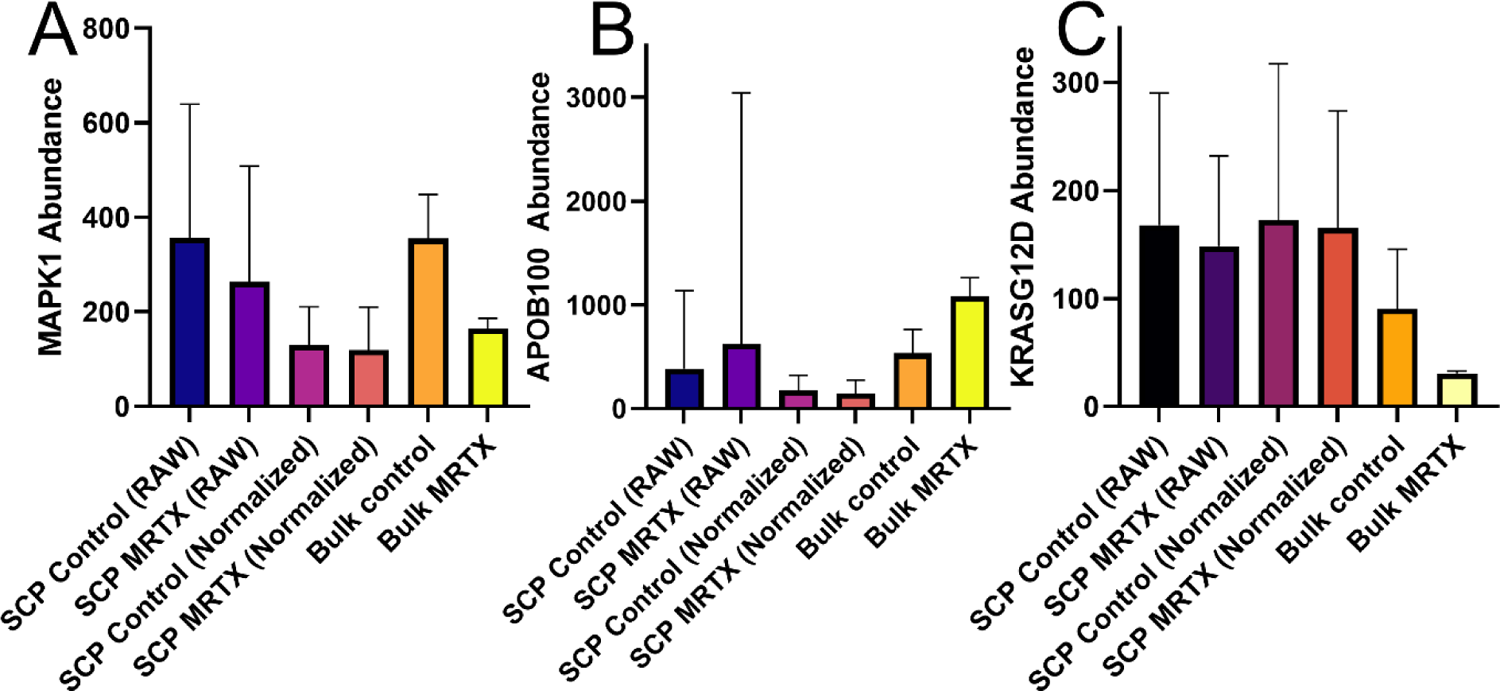
A demonstration of ratio suppression after protein normalization in SCP data, with non-normalized data (RAW) plotted against normalized and bulk cell proteomics findings. **A.** The abundance of MAPK1. **B.** The same analysis for Apolipoprotein B100. **C.** The abundance of the KRAS protein in PANC 0813 cells following treatment with MRTX1133.

Further spot checks of proteins observed as altered in the bulk proteomic data following treatment likewise found quantitative alterations suppressed by these typical normalization and scaling tools. Apolipoprotein B100 was a protein with substantially higher abundance than MAPK1 which followed the opposite trend, with a 2.03-fold increase following drug treatment in PANC 0813 bulk proteomics. Measurements of the mean of the original peptide mirror the differential regulation observed in the bulk proteomics experiment, but this observation is lost following normalization and scaling. (Figure 6B). The observation that data handling can reduce quantitative significance in SCP is not new to this study. A label free SCP work noted that imputation markedly reduced peptide differentials when applied.^35^ A recent reanalysis of those files further determined that recapitulation of bulk proteomics from SCP was convoluted by both standard normalization and imputation approaches.^36^ While a full analysis of the effects of commonly used tools for bulk proteomics on SCP data is beyond the scope of this manuscript, these results do suggest that new informatics should be embraced for single cell proteomics, and the results of traditional tools should be interpreted with caution.

### Grouping MRTX1133 treated cells by total KRAS abundance suggests subpopulations exist

As expected for a KRAS inhibitor, treatment with MRTX1133 leads to an overall decrease in KRAS protein abundance in the bulk proteomics data. Likewise, the mean of the non-normalized abundance of KRAS protein in single cells demonstrated a net decrease in protein abundance. The recapitulation of bulk proteomic findings is a useful confidence metric for SCP data interpretation, but it isn’t the goal. The real power in SCP is the ability to identify and evaluate subpopulations of cells with different phenotypes.^37^ The mean protein abundance of KRAS in untreated PANC 0813 cells was calculated at 167.8, while the same for treated cells was 124.1 (Figure 6C). Approximately 15.1% of MRTX1133 treated PANC 0813 cells were observed to have a KRAS protein abundance greater than the total population mean. These results suggested that multiple populations of cells may be present following drug treatment at this time point, with at least one population of cells where the drug reduced total KRAS expression and one where it did not. To further evaluate this subpopulation, treated cells with a non-zero measurement for KRAS protein expression were clustered for SimpliFi statistical analysis with the cells divided into two groups, those with above median KRAS protein abundance and those below that line. From this analysis, 62 analytes were determined as significantly differential (log 2 > 1, p-value < 0.05). The MAP kinase 4/6 pathway again appears differential in this experiment (R-HSA-5687128, p = 6.88 x 10^-8^). In addition, several proteosome subunits, namely 26S subunits 13 (PSMD13) and both Alpha type 3 and 5 (PSMA3, PSMA5) appear significantly higher in abundance in cells with higher mutant KRAS protein abundance. A complete list of differential proteins and statistical analysis is provided as **Supplemental File 3**. When evaluating proteins as differential by fold-change alone, proteins involved central metabolism and protein translation as well as proteosome degradation appear to be the highest in cells with above median KRAS abundance. Lower relative KRAS expression corresponds to increased levels of Ubiquitin 6, 10 and E2 all having 4-fold higher abundance in cells with low KRAS abundance. While full conclusions may be difficult to draw from a relatively small sample set, it is tempting to conclude that in cells where mutant KRAS protein has not been successfully inhibited at this dose and time point, the cells are proceeding with business as usual. In cells where inhibition has occurred effectively, the lack of the pro-survival cascades caused by the mutant protein has inhibited cell growth and metabolism and are beginning to lead to quiescence or apoptosis.

### In cases where drug treatment may result in large cell death single cell proteomics may not be possible

PANC 0203 and PANC 0403 have been recently described as two cell lines with extremely high sensitivity to MRTX1133.^5^ My results strongly support these claims as 10 nanomolar treatment for 48 hours resulted in both dramatic reductions in viability as well as alteration in cellular morphology as detected by fluorescence scatter. During single cell isolation in this study, viable cells were collected based on scatter using a simple live/dead cell stain. For both PANC 0203 and PANC 0403 cells it was not possible to collect 200 cells for the carrier channel for more than a few intended plates due to the reduced number of cells present. Sorting gates required adjustment for both these treated cell lines due to the level of difference in relative scatter compared to control (**Supplemental** Figure 2). In this case, the cells that were isolated appeared to be the minority of all cells leading to concerns regarding the value of data from these cells for this study. Repeated attempts to obtain single cell proteomics data on the PANC 0203 cell line have been unsuccessful even at levels of drug treatment significantly lower than previous reports for this cell line (data not shown). For treatment conditions where large amounts of cell death may be an outcome, careful titration of conditions may be necessary to obtain biologically meaningful cells for study. In some cases, this may not be possible at all.

### Limitations of this current study

Considerable limitations exist in this current work as described. Limitations in the metabolomics analysis include the lack of internal or external controls for the identification and relative quantification of the identified metabolites. All identifications were made through a combination of high resolution MS1 and MS/MS identifications against spectral library databases. As no tools have been fully implemented for the estimation and false discovery rates of metabolomics data today, library based matches should always be interpreted with caution. Single cell proteomics data comes with a myriad of limitations including that a data set of this size should be interpreted by a larger team than a single. While this work was intended to pressure test our current SCP workflows while evaluating a drug of particular interest to our program, the true goal of our work is to analyze time courses of drug treatments. While this will require significantly more time, cells and assistance to execute, the basic workflow appears sound to carry out these studies.

## Conclusions

In this work I describe the application of metabolomics, proteomics and single cell proteomics to the analysis of acute dosage of a promising new small molecule inhibitor. While MRTX1133 is currently in clinical trials and may not succeed, it is inevitably the basis of further compounds to target one of the most deleterious of all human cancer mutations.^38^ To the best of my knowledge, this is the first metabolomic or proteomic analysis of cells treated with this compound. Metabolomics demonstrates that the drug leads to a decrease in both ATP and NAD precursors in all cell lines. In addition, many amino acids appear to accumulate following drug treatment while some amino acids such as glutamine appear to be differential between cell lines in a manner that corresponds to the number of mutant copies of the gene in each cell line. However, this is the result of only four cancer cell lines and these results should be interpreted with caution. Proteomic analysis indicates the expected decrease in the MAPK pathway in all cell lines as well as an alteration in GTPase signaling and MHC expression. In general, results that should be expected based on previous results.^1,39^ To add further granularity to these measurements, I analyzed approximately 1,500 single cells from the PANC 0813 and ASPC-1 cell lines.

In addition, I divided the single cell study into two sections, one in which the carrier channel was a collection of presumably viable FACs sorted cells, and a second where the carrier was created from a bulk cell homogenate. These two methods for preparing the carrier channels imparted differences in the proteins that were observed. While many of the protein level differences between cells analyzed with the two carrier channel strategies are difficult to reason through, some are not. The presence of Caspases 6 and 8 were exclusively detected in single cells where a bulk cell lysate carrier channel was employed. This suggests that when FACs isolate viable cells as carrier channels, proteins from cells that are dead or dying are being sorted away with the unhealthy cells they belong to. In the case of drugs such as MRTX1133 where the goal is the death of cells harboring a specific mutation care must be taken to assess the correct population for each experimental goal.

Considerable effort today is being directed toward the improvement of the field of single cell proteomics. The majority of work to date has centered on developing new tools for cellular isolation, peptide recovery, and increased measurement sensitivity.^18,27,40,41^ However, a recent multi-institute collaborative effort has provided suggestions for the reporting of single cell proteomics data^42^ and informatics has continued to develop to address the unique challenges inherent in this emerging new field.^43^ My takeaway from this study is that equal effort may be necessary to develop ideal experimental designs for a successful SCP study on drug treatment models.

## Acknowledgements

I would like to thank Hannah Wilkins for assistance with PANC 0203 label free experiments and advice on cell culture and Dr. Alejandro Brenes for sharing unpublished information on TMTPro reagent optimization for minimizing interference. Ahmed Warshanna and Dr. Hao Zhang deserve thanks for staying late on the last day of the 2022 school year to help me get these cells isolated. Finally, I would like to thank Jim Palmieri and John Wilson for sharing an unpublished reanalysis of the data deposited with the preprint of this manuscript.

## Supporting information

The following supporting information is available free of charge at ACS website: http:/pubs.acs.org

**Supplemental Figure 1.** Proteins exclusively observed when a bulk cell homogenate sample is used as a carrier channel.

**Supplemental Figure 2.** FACs analysis demonstrating increased cell death in MRTX treated PANC 0203 cells.

**Supplemental File 1.** Metabolomic alteration in four MRTX1133 treated cell lines

**Supplemental File 2.** Proteomic report for all single cells in this study

**Supplemental File 3.** A summary of single cell proteomic observations of MRTX1133 treated cells with above median KRASG12D protein expression versus those with below median expression.

