## Supporting Information and Supplemental Figures for "Metabolomic, proteomic and single cell proteomic analysis of cancer cells treated with the KRAS^G12D^ inhibitor MRTX1133"

**Benjamin C. Orsburn^*^**

The Department of Pharmacology and Molecular Sciences

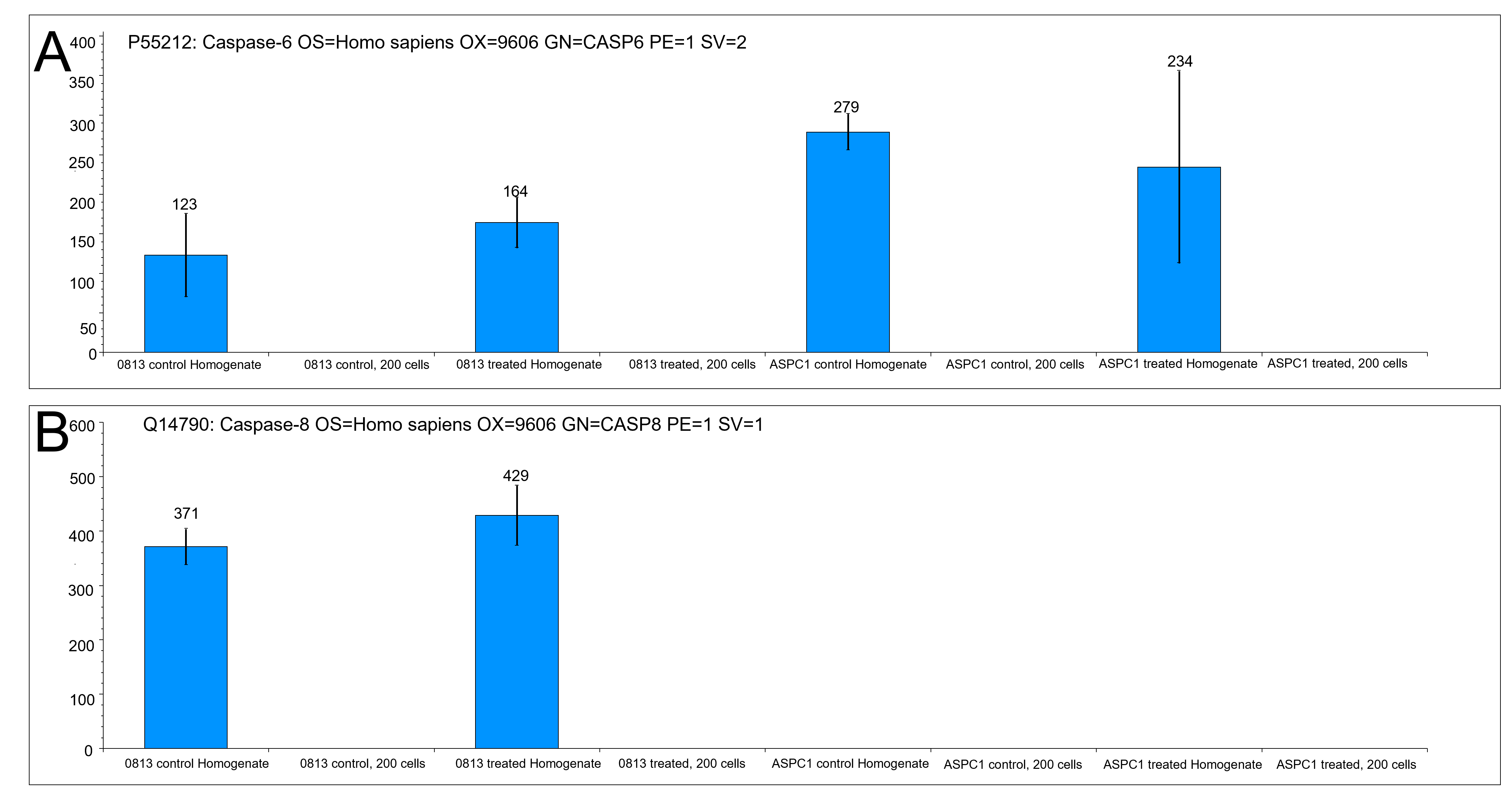

**Supplemental Figure 1. Proteins exclusively detected when a bulk cell homogenate is used for a carrier channel. A.** Caspase 6 which was detected in all homogenate experiments. **B.** Caspase 8 which was exclusively detected in PANC 0813.

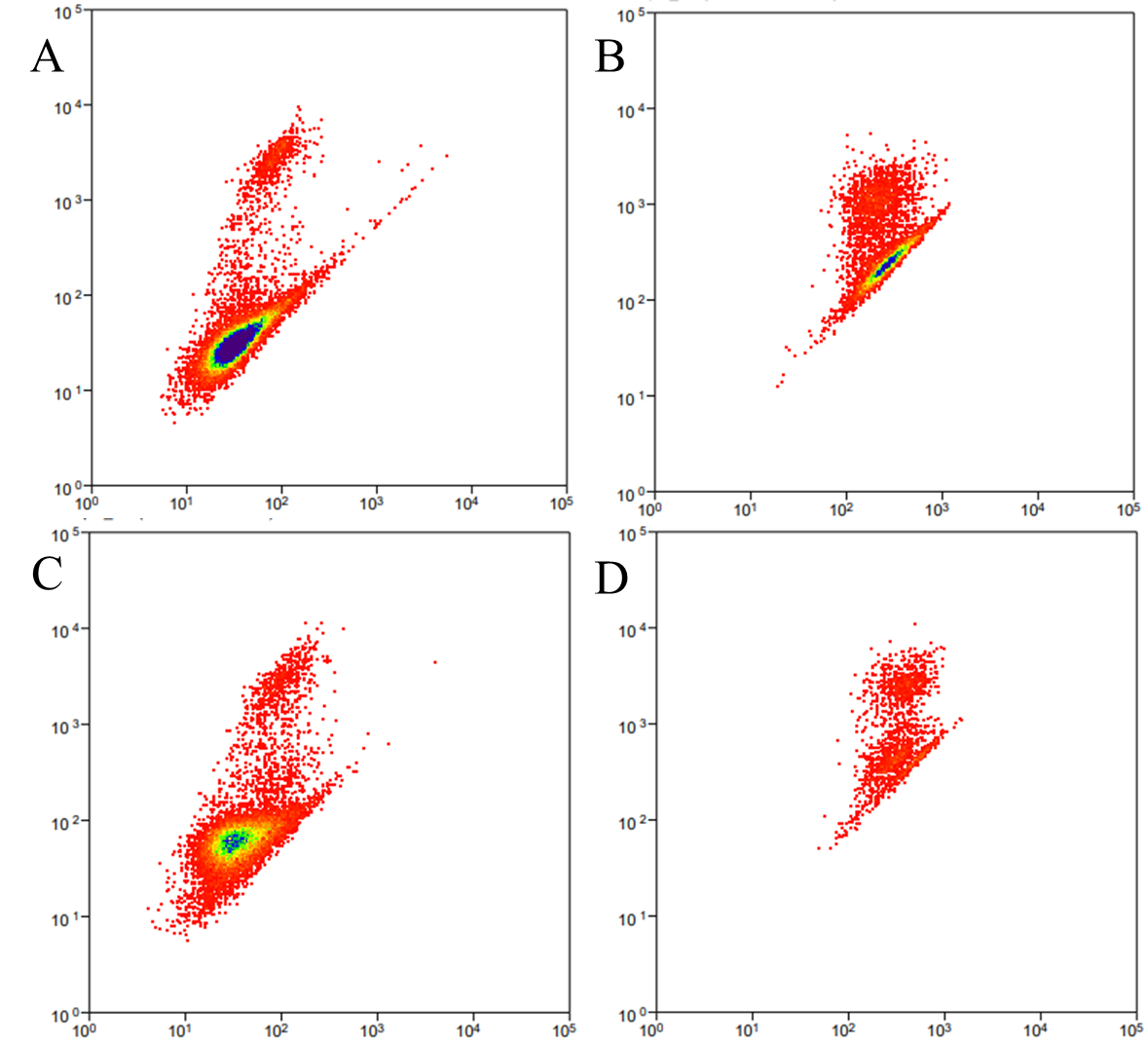

**Supplemental Figure 2. Fluorescence output data for two KRASG12 mutant cell lines following acute treatment with MRTX1133.** **A**. ASPC-1 control cells. **B**. ASPC-1 cells treated with MRTX1133 **C.** PANC 0203 control cells. **D.** PANC 0203 cells treated with MRTX1133.
